# AI-Based Antibody Design Targeting Recent H5N1 Avian Influenza Strains

**DOI:** 10.1101/2025.04.24.650061

**Authors:** Nicholas Santolla, Colby T. Ford

**Affiliations:** University of North Carolina at Charlotte, School of Data Science, Charlotte, NC, USA; University of North Carolina at Charlotte, Center for Computational Intelligence to Predict Health and Environmental Risks (CIPHER), Charlotte, NC, USA; University of North Carolina at Charlotte, Department of Bioinformatics and Genomics, Charlotte, NC, USA; Tuple LLC, Charlotte, NC, USA

**Keywords:** Avian Influenza, Antibodies, Protein Modeling, Docking, AI

## Abstract

In 2025 alone, H5N1 avian influenza is responsible for thousands of infections across various animal species, including avian and mammalian livestock such as chickens and cows, and poses a threat to human health due to avian-to-mammalian transmission. There have been 70 human cases of H5N1 influenza in the United States since April 2024 and, as shown in recent studies, our current antibody defenses are waning. Thus, it is imperative to discover new therapeutics in the fight against more recent strains of the virus.

In this study, we present the *Frankies* framework for automated antibody diffusion and assessment. This pipeline was used to automate the generation of 30 novel anti-HA1 Fv antibody fragment sequences, fold them into 3-dimensional structures, and then dock against a recent H5N1 HA1 antigen structure for binding evaluation. Here we show the utility of artificial intelligence in the discovery of novel antibodies against specific H5N1 strains of interest, which bind similarly to known therapeutic and elicited antibodies.

## Introduction

Highly pathogenic avian influenza A (H5N1) remains a persistent global health threat due to its zoonotic potential and historically high mortality rate in humans. Despite sporadic outbreaks and ongoing disease surveillance by the World Health Organization, United States Department of Agriculture, and US Centers for Disease Control, the continual antigenic evolution of the hemagglutinin (HA) glycoprotein, particularly within the HA1 subunit, presents significant challenges to the development of broadly neutralizing therapeutics. The HA1 domain is critical for receptor binding and immune recognition, making it a prime target for antibody-based intervention (1).

To address the urgent need for rapid, scalable design of high-affinity antibodies against emerging H5N1 variants, we introduce *Frankies*, a computational pipeline that integrates state-of-the-art generative artificial intelligence (AI) modeling, protein structure prediction, and docking simulations. This pipeline begins by generating novel Fv antibody fragment sequences via conditional protein diffusion using EvoDiff, leveraging a curated set of reference sequences from the Therapeutic Structural Antibody Database (Thera-SAbDab). The resulting sequences were folded into 3-dimensional structures using structure prediction models such as AlphaFold3 or ESMFold v3, which have demonstrated near-experimental accuracy in modeling antibody-antigen interfaces (2, 3). Finally, we employ HADDOCK3 to perform rigid and flexible docking of the modeled Fv regions to the HA1 target, enabling the assessment of binding orientation, stability, and epitope accessibility.

By systematically combining data-guided AI generation with physics-based modeling, *Frankies* represents a promising step toward automated antibody discovery and optimization. In this study, we apply the *Frankies* pipeline to a recent H5N1 HA1 antigen and characterize a set of *de novo* antibody candidates with favorable structural and biophysical properties. This work lays the foundation for future *in vitro* empirical validation and paves the way for accelerated response to future influenza outbreaks.

## Results

### Pipeline Performance

The protein generation pipeline successfully produced 30 novel anti-HA1 Fv antibody fragment sequences, with a 100% success rate in AbNumber sequence validation. In our systematic tests, the reliability of Fv antibody sequence generation was consistently greater than 90%. The complete set of 30 experimental runs took 11 hours and 34 minutes to complete in serial, averaging 21.13 minutes per run. In a distributed computing context, this walltime would scale linearly with *∼*20 minutes per pipeline run. Individual component processing times were distributed as follows: approximately 5 minutes for EvoDiff operations, less than 1 minute for structure predictions with ESM3 (or around 20 minutes with AlphaFold3), 16 minutes for HAD-DOCK3 computations, and less than 1 minute for the output report generation, shown in Figure 1.

**Fig. 1.**
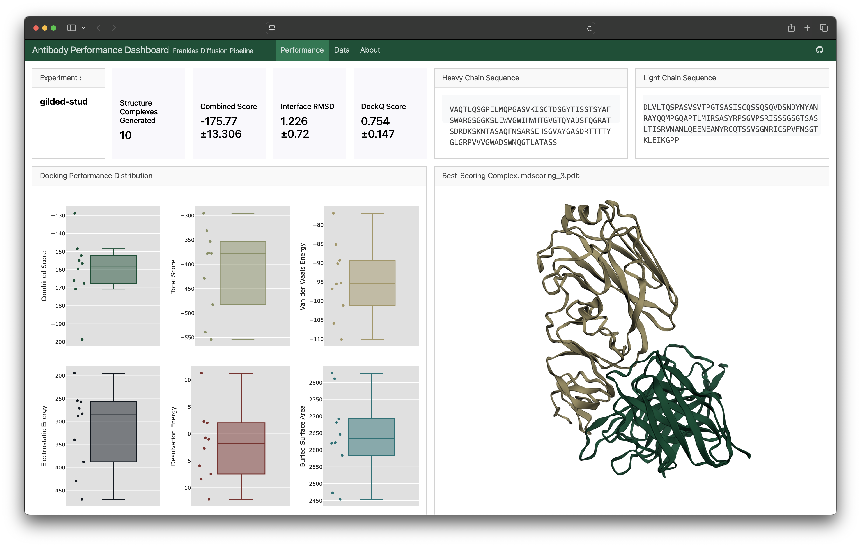
*Frankies* output dashboard report. Generated with Quarto.

### Generated Fv Sequences

The 30 diffused Fv heavy and light chain sequences are listed in Table 1. Note that all diffused sequences are a defined length, with heavy and light chains having 132 and 125 amino acids, respectively. This is controlled by the diffusion step, which defines a maximum sequence length to be conditionally generated. The top 10 candidates, by lowest HADDOCK score, are shown in Figure 2. Note the epitope diversity given the random surface residue detection for unbiased binding site selection.

**Table 1.**
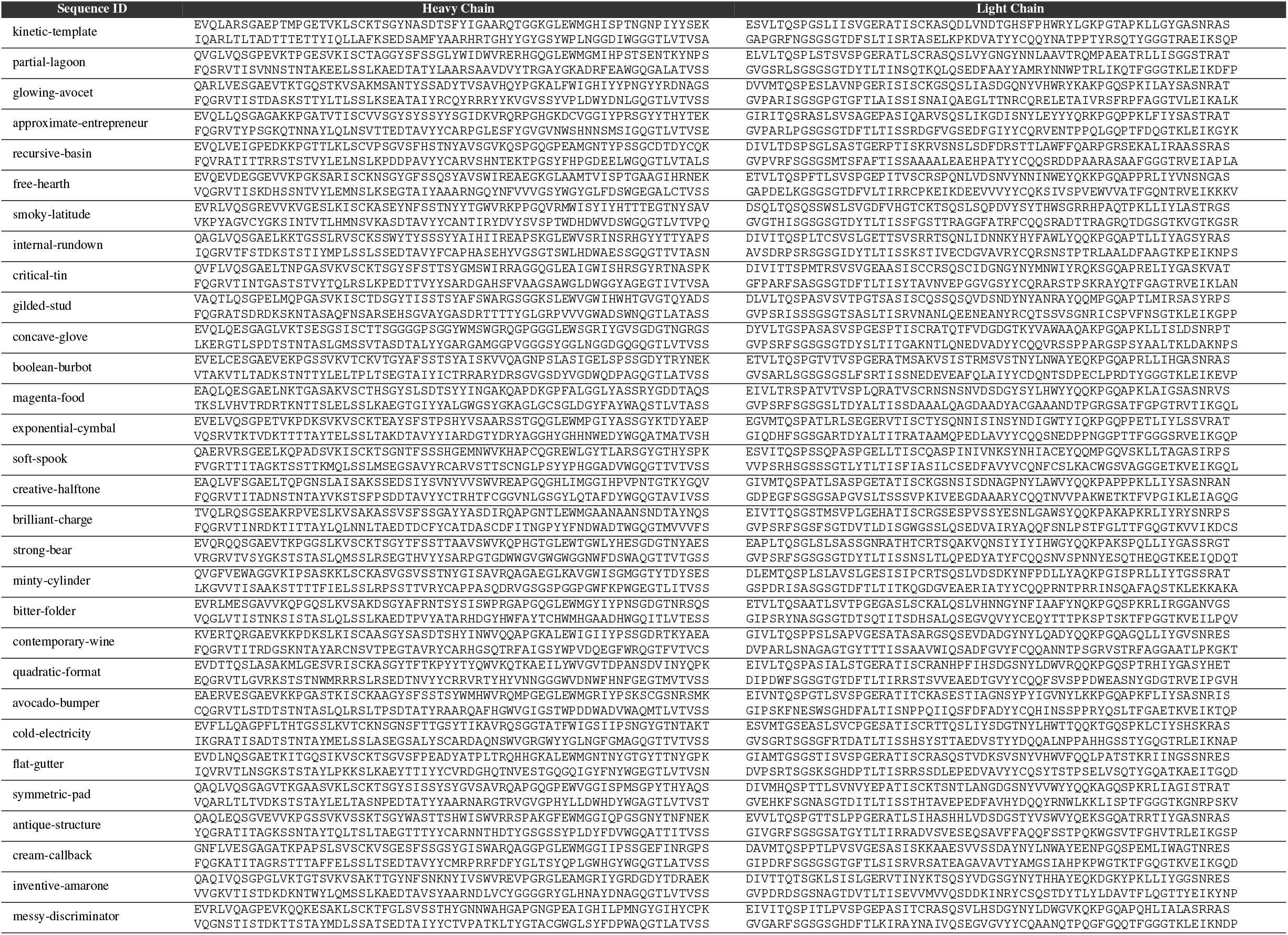
The 30 Fv heavy and light chain sequences diffused and evaluated in this study.

**Fig. 2.**
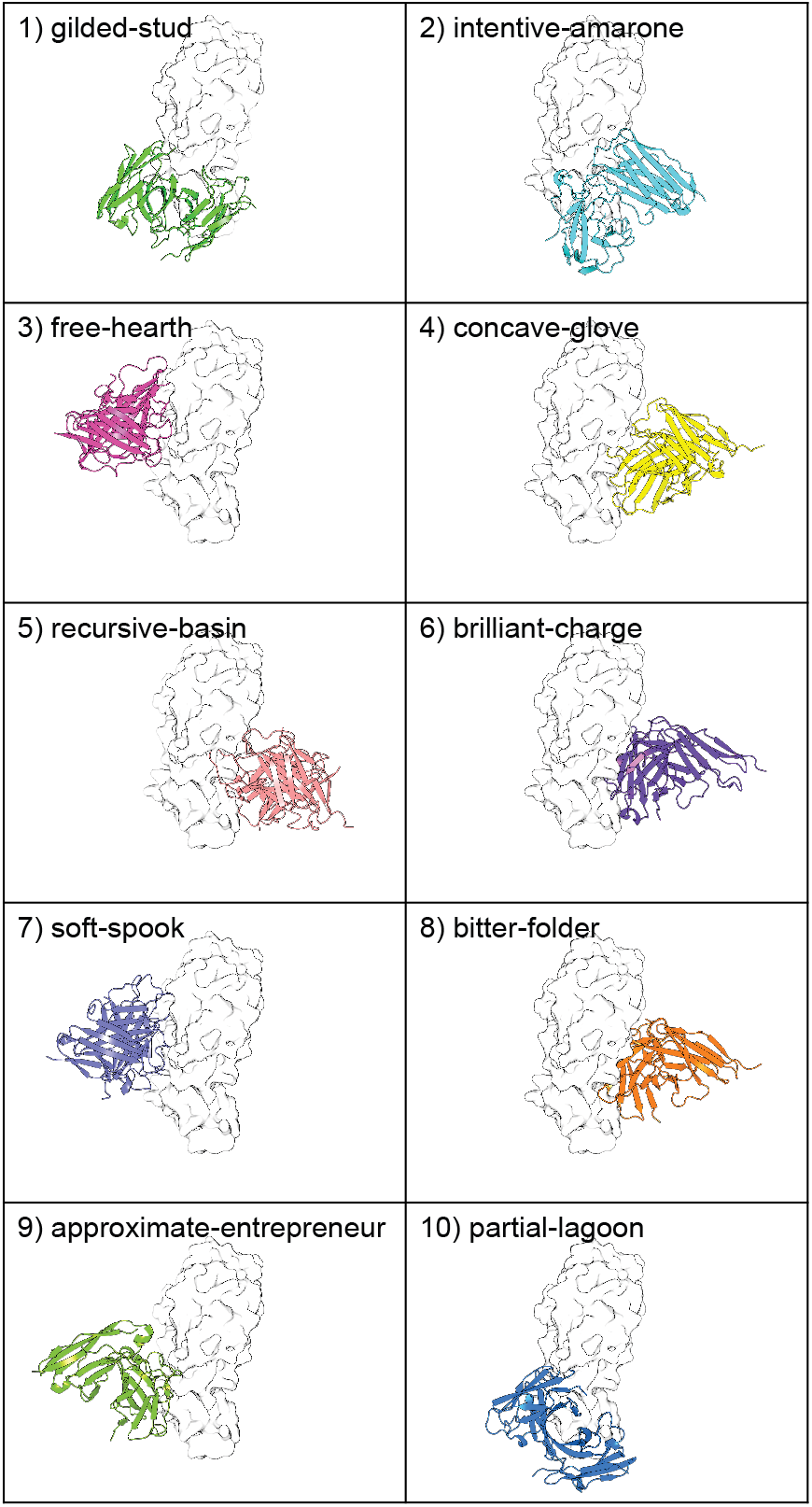
Top 10 diffused Fv candidates by HADDOCK score. EPI3009174 HA1 is shown as a white surface model and the diffused Fv structure is shown in a colorized cartoon style.

### Epitope Variation

In the *Frankies* pipeline, users have the option of defining active residues on the antigen to guide the docking process. In this study, however, we allowed for the diffused antibodies to bind to any surface residue on the antigen structure, reducing the bias in the binding process. This produced an interesting pattern of particular residues commonly forming polar contacts with the antibody CDR loops. See Figure 3.

**Fig. 3.**
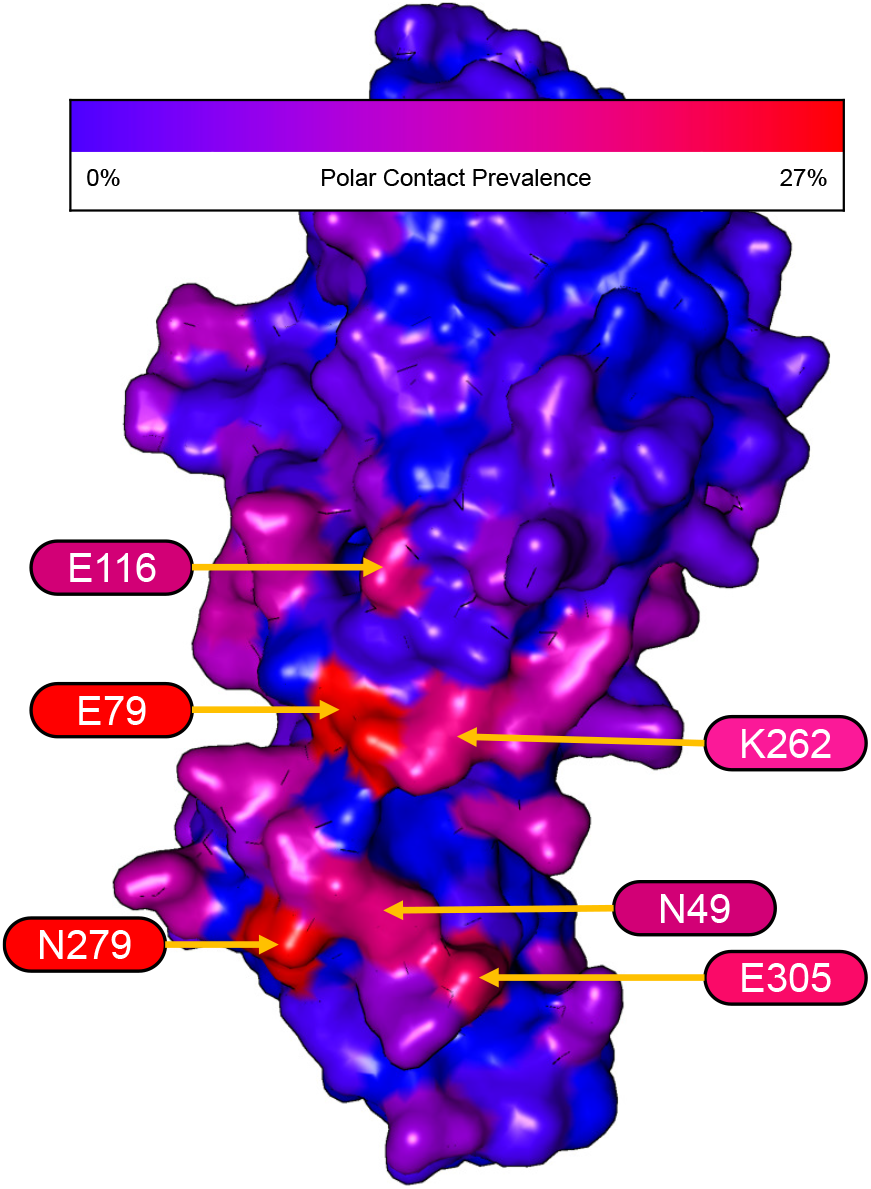
Structure heatmap showing the prevalence of polar contacts made between the diffused antibodies and the HA1 antigen. Annotated residues indicate those that are interfaced *≥* 20% of cases across the 30 diffused antibodies in this study.

#### Structure Biochemistry

Further evaluating the biochemistry of the diffused Fv structures shows consistent desirable traits across various metrics.

Evaluating protein stability, desirable solvation metrics were predicted using FoldX. As shown in Figure 4A, the average total energy predicted by FoldX was -121.9 kcal/mol, ranging from -144.8 to -60.7 kcal/mol. Also, favorable polar solvation and hydrophobic solvation was predicted, indicating desired polar interactions with the solvent along with proper burial of hydrophic residues in the Fv structures. Futhermore, the predicted hydrophic solvation indicates a low propensity for aggregation of the proteins *in vivo* (4). See Figures 4B and 4C.

**Fig. 4.**
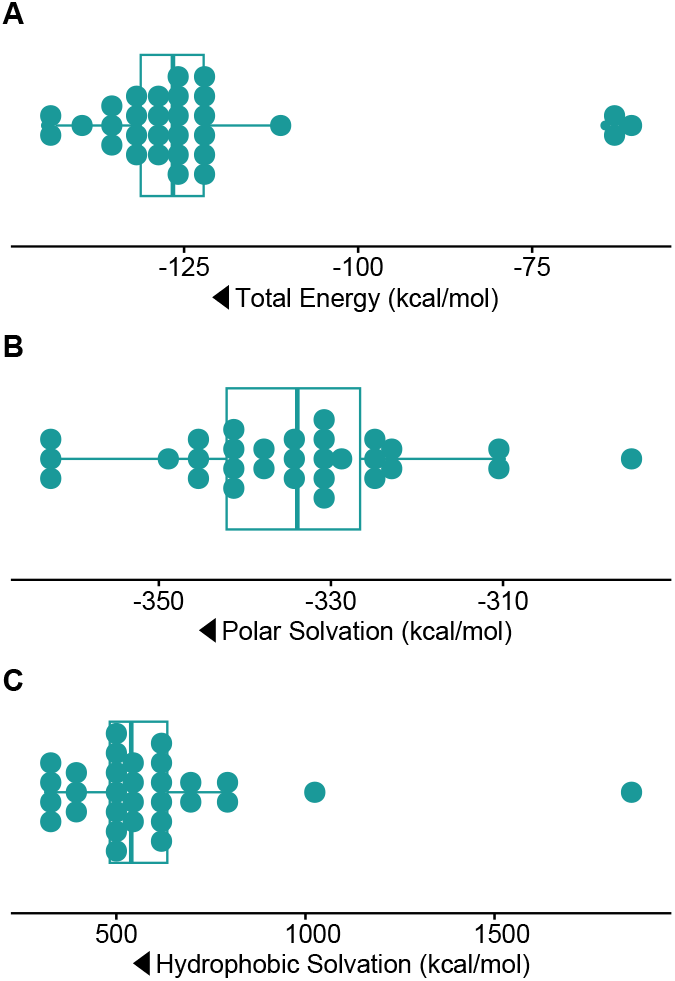
Boxplots depicting the distribution in solvation metrics. A) Total energy - More negative values indicate better overall stability. B) Polar solvation - Lower values indicate favorable polar interactions with solvent. C) Hydrophobic solvation - lower values indicate better burial of hydrophobic residues (favorable for folding) and high values might suggest exposed hydrophobic surfaces (i.e., risk of aggregation or low solubility). Arrows indicate the “better” direction of each metric.

Regarding the humanness of the diffused heavy and light chain Fv sequences, we see a bimodal distribution where about half of the chain sequences are predicted to be human (whereas the remaining are of hybrid-to mostly murine-level of composition), shown in Figure 5.

**Fig. 5.**
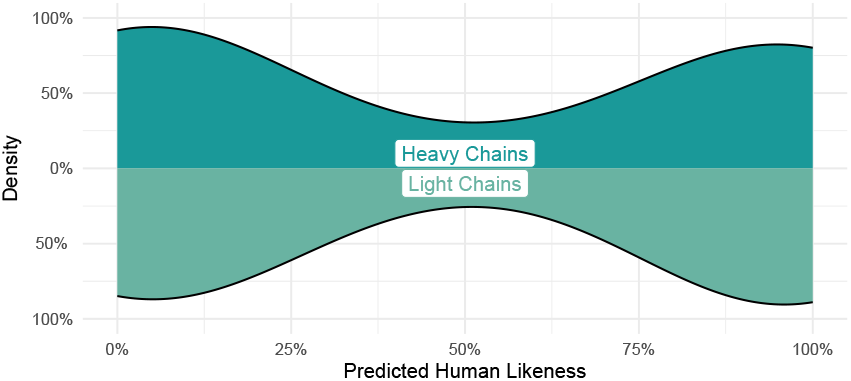
Density plots showing distribution in the predicted humanness of the diffused Fv heavy and light chain sequences.

Note that none of the sequences in this study have been humanized. Rather, the diffusion process simply generated sequences on a spectrum between human and murine, based on the input set of reference antibodies, preferring either extrema rather than creating hybrid sequences. Thus, the humanness of these diffused antibodies can be improved using standard humanization tools/processes. The average humanness probability for the heavy and light chains of the top 10 performing antibodies, shown in Figure 2, was 44.7% and 62.2%, respectively.

### Sequence Analyses

Note the sequence diversity depicted in the logos in Figures 6a and 6b. While natural heavy chain sequences often start with EVQ or QVQ, there is additional sequence diversity in position 3 with the introduction of arginine (R), phenylalanine (F), glycine (G), and aspartate (D) amino acids, which are atypical here. However, the conditional sampling shows a consistent selection of glycine at the 6th position and VTSS at the end of the VH sequence, which are very common in natural antibodies. For the light chains, the common starting sequences of EIV and DIV were seen in the diffused sequences along with expected sampling of methionine (M) or leucine (L) at position 4, and EIK toward the end of the VL sequence (positions 120-122) (5). Note that all of these sequences were successfully numbered using the Chothia numbering scheme (6).

**Fig. 6.**
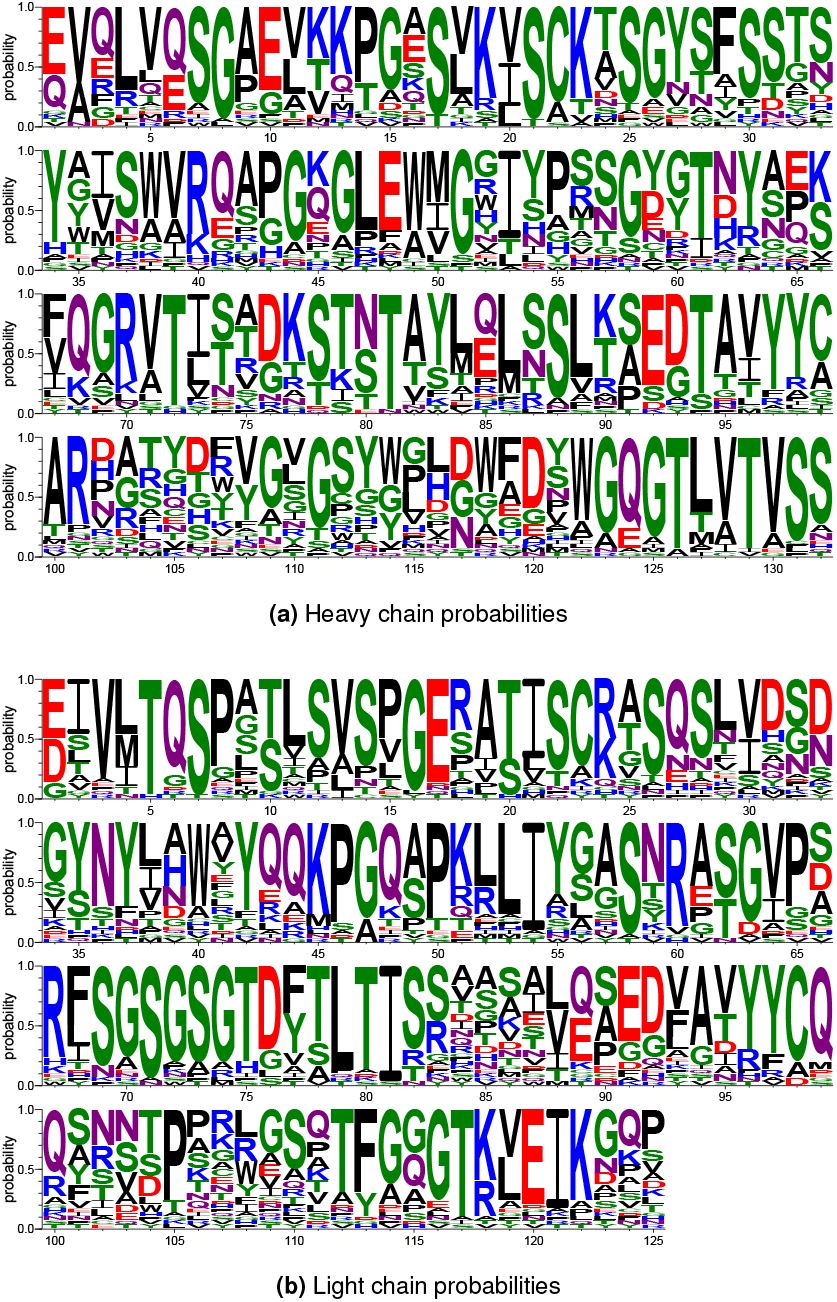
Position-level amino acid probabilities of the diffused Fv chain sequences. Created with WebLogo (7).

Comparing the sequences of the diffused and reference antibodies shows a 47.24% (*±*0.06%) identity in the heavy chains and a 52.37% (*±*0.08%) identity in the light chains. This is within an expected range given that the generated sequences were diffused by sampling an input distribution and that all of the sequences folded into the desired Fv antibody shape. Sequence identity distributions and more detailed pairwise identity comparisons are shown in Supplementary Figure 1.

#### Folding Performance

Of the 30 diffused pairs or Fv sequences presented in this study, all folded with an average Predicted Local Distance Difference Test (pLDDT) confidence >0.5. As shown in Figure 7, most sequences folded with a mean and median pLDDT >0.75, indicating a highly confident structure prediction.

**Fig. 7.**
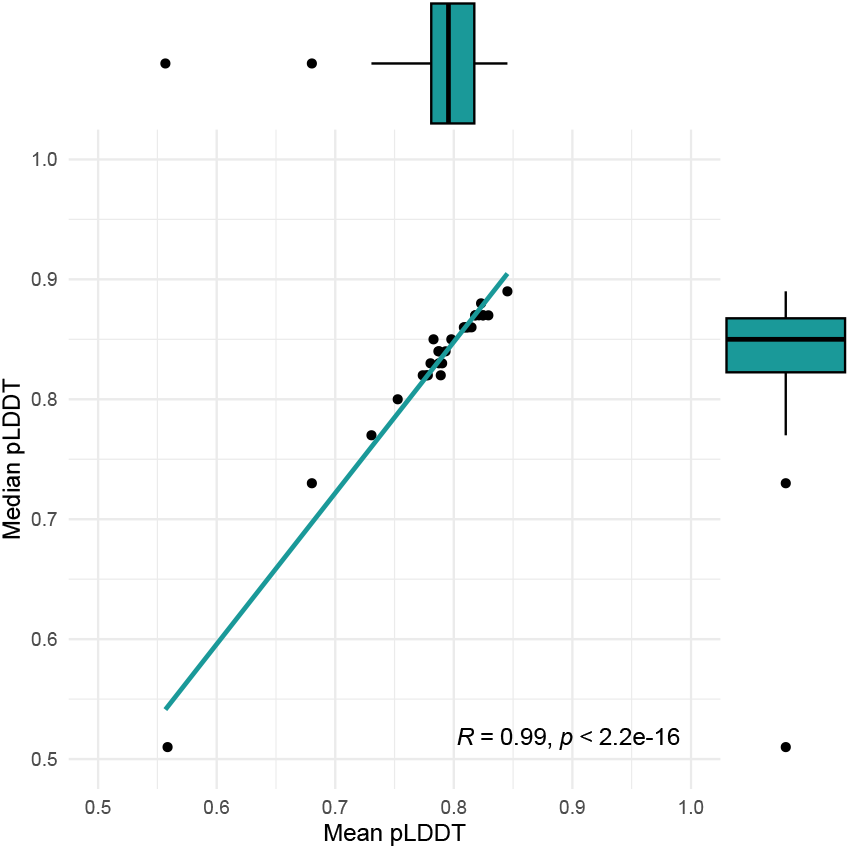
Scatterplot of mean versus median pLDDT across the 30 diffused Fv structures folded with ESM3. Marginal boxplots show the distribution of the mean and median pLDDT values.

This is expected given that each diffused Fv sequence was numbered through the Chothia numbering system as a quality control step in the *Frankies* pipeline before advancing through to the evaluation steps.

### Comparison to Existing Antibodies

The *Frankies* pipeline consistently generated Fv sequences and structures that bound well to the HA1 epitope.

#### Binding Performance

For comparison purposes, as shown in Figure 8, the binding performance of the 11 antibodies used in Ford et al. 2025 that were bound to H5N1 isolate EPI3009174 were selected as a reference. Then, these were compared to the 30 Fv structures generated in this study, which were docked against the same isolate antigen structure. The Van der Waals energy is significantly better (more negative, statistically) in the diffused antibodies, indicating stronger non-bonded atom-atom interactions. This could imply tighter packing or improved interface complementarity in the diffused antibodies.

**Fig. 8.**
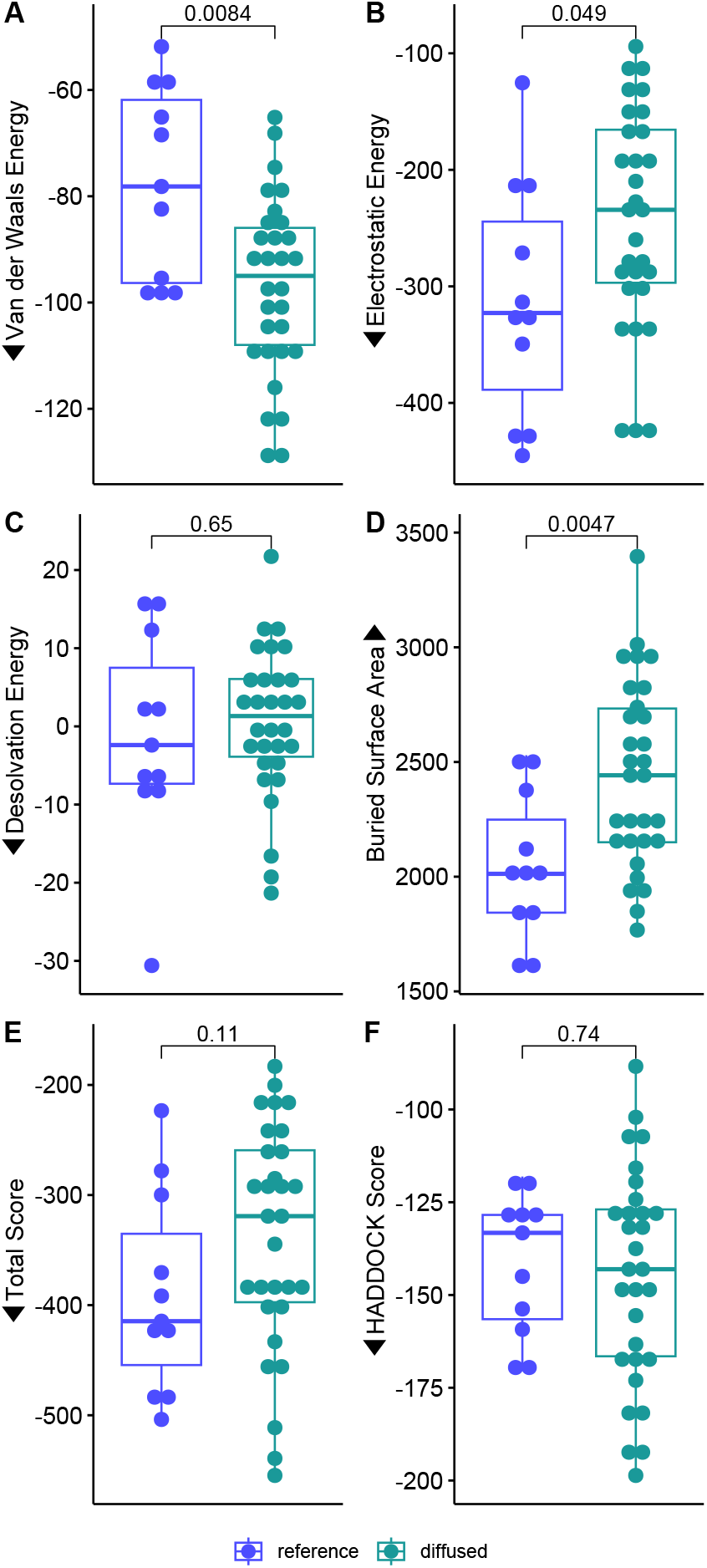
Comparison of the docking metrics between reference and diffused antibodies. Pairwise comparisons are shown as p-values from the Wilcoxon signed-rank test. For all metrics except buried surface area and desolvation energy, lower is likely indicative of better binding, indicated by arrows.

In contrast, the reference antibody set displayed better electrostatic energy and buried surface area, though there are multiple examples of diffused antibodies that are within the same range. A larger buried surface often correlates with stronger binding and greater stability. In this case, the diffusion process may be generating antibodies with slightly reduced interfacial engagement.

However, the desolvation energy, total score (electrostatic + Van der Waals energies), and HADDOCK score were quite similar overall and showed no statistical difference according to the Wilcoxon signed-rank test at the *α* = 0.05 level.

Thus, these metrics show that this pipeline was able to consistently generate antibody sequences with similar performance to therapeutically-derived or elicited antibodies.

## Methods

The *Frankies* pipelines was designed as an automated workflow for producing antibody candidates and predicting their binding affinity at-scale. This pipeline is written in Snakemake (9), which provides a reproducible framework for running all of the steps mentioned below.

Inside the Snakemake pipeline, various steps are run as Python or Shell scripts, while other complex tools are executed using Docker Containers. The modularity of the pipeline is designed such that each run produces a single antibody candidate, tagged with a randomly generated experiment name, along with an evaluation of its performance against a given target. Thus, the *Frankies* pipeline can be run *n* times, without modifying the initial configuration, to produce *n* unique Fv candidates. The overall workflow is shown in Figure 9.

**Fig. 9.**
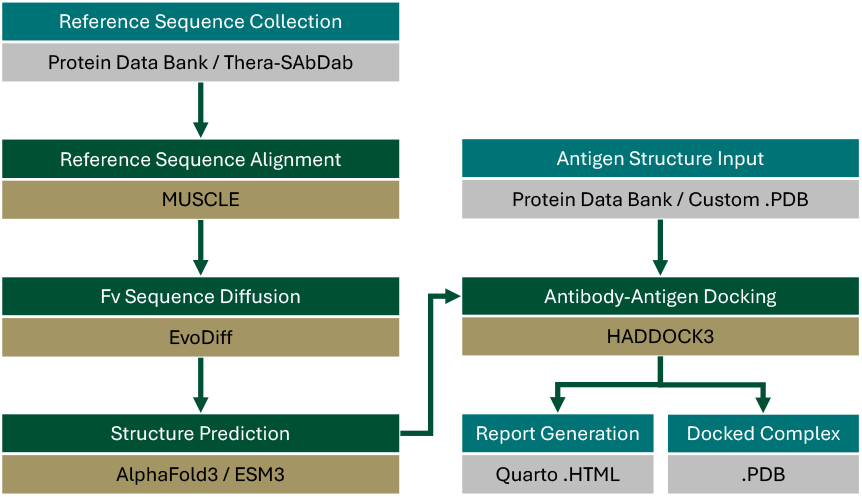
*Frankies* pipeline workflow steps

### Reference Dataset Preparation

To guide conditional sequence generation, we curated a reference set of HA1-targeting Fv sequences from the Therapeutic Structural Antibody Database (Thera-SAbDab) combined with additional antibodies used in Ford et al. 2025 listed in Table 2.

**Table 2.**
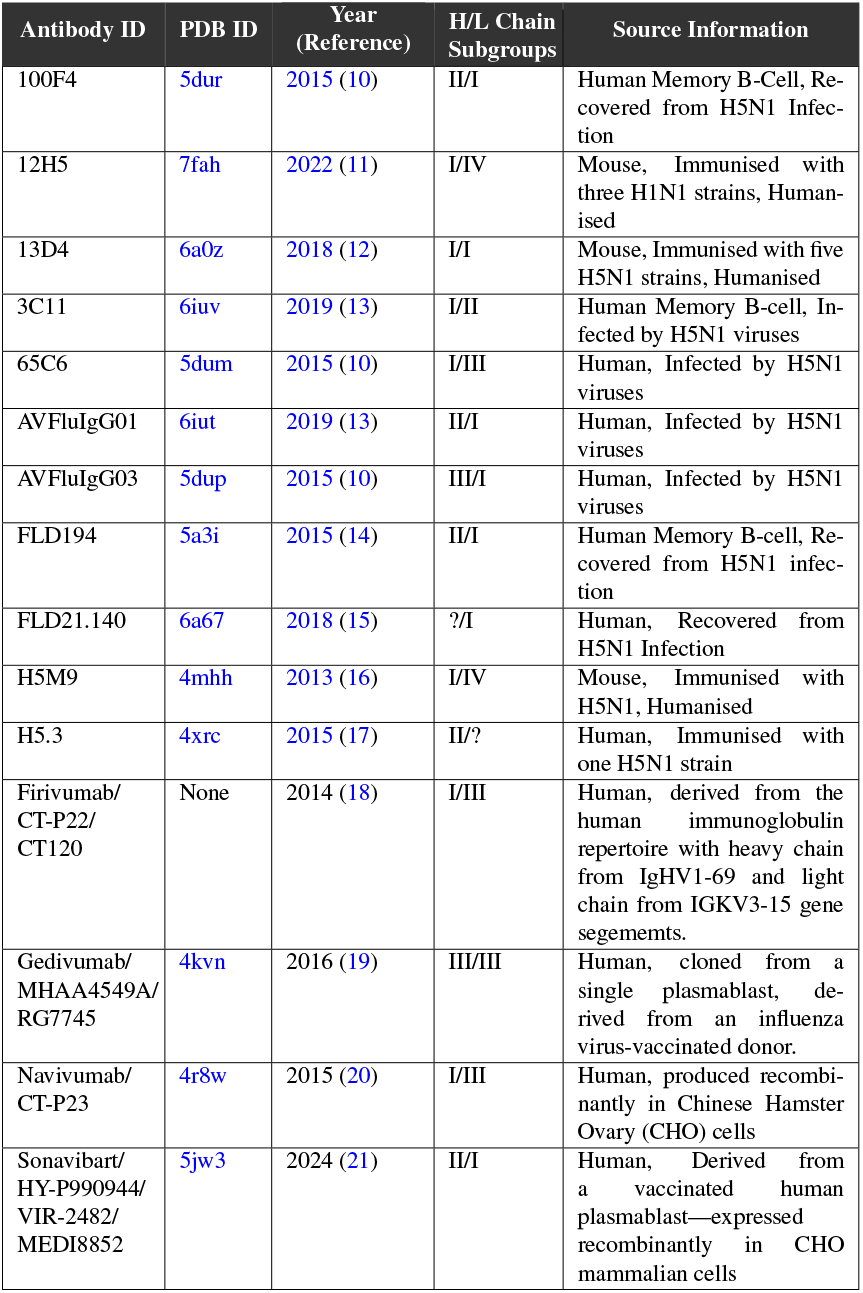
Reference HA1-neutralising antibodies from TheraSAbDab and those used in Ford et al. 2025.

Sequences were trimmed to retain only the Fv portion of the antibody chains and were then aligned using MUSCLE v3.8.425 (22). This produced separate input .a3m alignment files for heavy chain and light chain sequences.

### Conditional Sequence Generation

We used EvoDiff’s MSA_OA_DM_MAXSUB model for conditional sequence generation (23). The conditioned sequence generation sampled from the curated reference sequence files using the ‘Max-Hamming’ distance to produce novel, HA1-targeted Fv chain sequences. EvoDiff operates as a protein diffusion model trained on general sequence databases and can be guided by user-provided templates or alignments of desired reference sequences in .a3m format.

Thirty Fv candidate sequences were generated and filtered for naturalness using AbNumber, a Python wrapper for ANARCI (24). This attempted to number the diffused sequences with the Chothia numbering system (6) and, if it failed, the diffusion process would restart generate another sequence. This was performed to help ensure that the diffused sequences were as antibody-like as possible, lending to more confident subsequent folding and improve future protein synthesis capabilities.

Sequences were analyzed to predict their overall stability, including polar and hydrophobic solubility using FoldX v5.1 (25) using the ‘Stability’ command. This provided a variety of energy metrics for each diffused Fv structure, all of which are reported in the supplementary GitHub link.

Using Humatch’s ‘classify’ functionality, each sequence’s “humanness” was also evaluated, which reports a predicted human probability percentage for a given input heavy and light chain sequence pair (26).

Also, sequence identity of each diffused sequence was compared to the set of reference sequences using the Biostrings v2.66.0 library in R v4.2.2 (27).

### Structure Prediction

Diffused heavy and light chain sequences were folded using either AlphaFold3 (2) or ESM3 (3). AlphaFold3 may be desirable for groups wishing to perform purely local folding. While AlphaFold3 provides highly accurate structure predictions, it requires the user to download and store >600GB of reference databases and model weights to run. Plus, the multimer prediction of the heavy and light sequences together take approximately 15 minutes on a GPU.

Alternatively, Evolutionary Scale’s ESM package and esm3-medium-multimer-2024-09 model allow for API-based multimer structure prediction without any reference databases. This package returns the predicted structure in a few seconds.

Both AlphaFold3 and ESM3 consistently produce reliable antibody structures and thus we provide support for both models.

Complementarity-determining region (CDR) loops were detected using ANARCI and residues belonging to the CDR loop structures were selected as “active residues” in the subsequent docking process.

### HA1 Antigen Preparation

A reference target antigen structure—HA1 subunit of H5N1 hemagglutinin—was obtained from the Protein Data Bank (PDB ID: 2VIR). This structure was used to test and validate the *Frankies* pipeline.

Then, the HA1 structure of a more recent isolate EPI3009174 was folded and used for subsequent novel antigen docking. Isolate EPI3009174 was collected in from a 9-year-old Cambodian male patient who passed away in 2024. This isolate was previously analyzed in Ford et al. 2025 against 11 reference antibodies.

For docking, the residue numbers in the antigen structure were incremented +1000, to avoid overlapping numbers with the antibody structure, and all residues were assigned to chain B.

Surface residues are automatically detected on the antigen structure and a 25% random subset of the surface residues was selected as “active residues”.

### Antibody-Antigen Docking

Fv structures were docked to the HA1 antigen using HADDOCK3 (28, 29) following the methodologies shown our recent previous studies (30– 32). Docking configuration files were generated using an antibody-antigen docking template from the Bonvin Lab (33). These configuration files, along with the other required docking files, were generated in a docking preparatory step in the *Frankies* pipeline.

Docking was performed in a multi-stage protocol including rigid-body energy minimization, semi-flexible refinement, and explicit (water-based) solvent modeling. Docked complexes were scored using HADDOCK’s built-in scoring function and further evaluated using the other biochemical/biophysical binding metrics listen below. The template for the HADDOCK protocol is available in the GitHub repository.

- Van der Waals intermolecular energy (*vdw*) in kcal/mol
- Electrostatic intermolecular energy (*elec*) in kcal/mol
- Desolvation energy (*desolv*) in kcal/mol
- Restraints violation energy (*air*) in arbitrary units
- Buried surface area (*bsa*) in Å2
- Total energy (*total*): 1.0*vdw* + 1.0*elec* in kcal/mol
- HADDOCK score: 1.0*vdw* + 0.2*elec* + 1.0*desolv* + 0.1*air*

### Candidate Ranking, Reporting, and Visualization

The “best” complex is selected from the output of complexes based on the docking conformation with the best (lowest) HADDOCK score of the best cluster of complexes.

This best cluster and the best scoring complexes are reported in a rendered Quarto dashboard as the final step of the *Frankies* pipeline (34). This report shows the distribution of the various binding metrics of all complexes in the best cluster. This also renders the 3D structure of the best model as an interactive object using Py3Dmol, a Python wrapper for 3Dmol.js (35). A screenshot of the dashboard is shown in Figure 1.

## Discussion

As H5N1 continues to evolve, it is imperative that therapeutic advances continue to better target modern influenza clades of interest (e.g., 2.3.4.4b (36)) and therefore reduce the risk of mortality and morbidity in humans. As of late 2024, there are over 50 licensed H5 vaccine candidates (37), though many of these are based on significantly older strains. For example, in the United States, the 3 licensed vaccines are from 2007, 2013 and 2020 (38).

Some vaccine candidates are currently being developed using mRNA technologies on newer strains in the U.S. by Moderna (mRNA-1018) (39) and Arcturus Therapeutics (ARCT-2304) (40). These were shown to be effective in animal trials and are currently in Phase I/II clinical trials. Also, therapeutic antibodies are being developed that target H5 HA1 from clade 2.3.4.4b. Multiple promising candidates were generated through hybridoma technologies, as reported in a recent preprint by Alzua et al. 2025.

AI-based antibody discovery is already well underway with a few candidates having moved into clinical testing, including anti-TL1A antibody by Absci (ABS-101), (42–44). Such previous studies generated antibody sequences or structures using large-language or diffusion models such as RFdiffusion (45), AntiBARTy (46), and Abdiffuser (47) and then optimized the candidates with a lab-in-the-loop iterative process.

The *Frankies* pipeline presented in this study offers a streamlined and modular approach for the design of *de novo* antibody candidates against rapidly evolving viral targets such as H5N1 influenza hemagglutinin. By integrating generative protein diffusion, structure prediction, and flexible molecular docking, *Frankies* enables end-to-end discovery of Fv candidates with structural and biophysical characteristics that support high-affinity binding to the desired antigen.

A key innovation in *Frankies* lies in its conditional generative architecture. By seeding EvoDiff with therapeutically validated antibody sequences from Thera-SAbDab, we impose domain-specific constraints that preserve critical structural motifs while enabling conditional exploration of novel sequence space. This balances diversity with developability, reducing the likelihood of generating non-functional or unstable designs. Furthermore, our use of AN-ARCI and other quality control steps help to ensure that only the most promising candidates advance through the pipeline.

The application of *Frankies* to the H5N1 HA1 domain highlights the feasibility of using AI-driven approaches for pandemic preparedness. HA1 remains a challenging target due to its rapid antigenic drift and glycan shielding. Nonetheless, several *Frankies*-designed Fv candidates exhibited strong binding affinity scores and favorable interface properties, suggesting potential for neutralization. Importantly, these designs were generated *in silico* within minutes, underscoring the value of generative pipelines for rapid therapeutic prototyping.

Despite its strengths, *Frankies* has some limitations. The current conditional sequence generation can be improved in the future by implementing structure-aware diffusion (i.e., diffusing the CDR loops while a reference antibody is bound at the epitope site on the antigen). Also, better flexibility in CDR loop lengths is necessary.

While our predictions showcase the consistency of the Fv sequence generation, experimental validation will be crucial to confirm the binding specificity, affinity, and neutralization potency of the proposed candidates. Plus, the Fv sequences generated will need to be expanded to include the other parts of the antibody (the rest of the Fab region, the hinge, and the Fc region).

Today, the pipeline utilizes Docker to run the diffusion, folding, and docking steps, enabling users to run these processes without complex dependency installation in their local environment. While this is useful for cloud-based scalability, we will also implement the ability to choose the containerization engine. For example, Apptainer/Singularity are more common in on-premises high-performance computing (HPC) environments.

The current docking approach does not account for glycosylation, which plays a significant role in HA1 surface shielding and antibody accessibility. While HADDOCK3 provides valuable binding predictions, it lacks the full thermodynamic and kinetic accuracy of more computationally intensive free energy calculations. Thus, future improvements to the pipeline will include the integration of a molecular dynamics step, such as OpenMM (48), to model the trajectory of the antibody-antigen complex and to predict the stability of the interaction.

Future directions include fine-tuning EvoDiff on antibody–antigen co-evolution datasets, and benchmarking against existing broadly neutralizing antibodies targeting H5N1. Additionally, the modular nature of *Frankies* allows for straightforward extension to other antigens, including other influenza subtypes, entirely different viral families, or even targets in other diseases (such as in oncology, as shown in Ford 2024).

In conclusion, *Frankies* demonstrates how recent advances in protein generative modeling and structure prediction can be combined into an automated and scalable pipeline for therapeutic antibody design. As AI tools continue to mature, we anticipate that pipelines like *Frankies* will become central components of the nextgeneration biosecurity and drug discovery infrastructure for pandemic preparedness and beyond.

## Contributors

Authors NS and CTF developed the *Frankies* pipeline. Author CTF performed data curation of the anti-HA1 antibody Fv sequences and their multiple sequence alignment. Author NS performed the docking experiments and statistical analyses. CTF and NS generated all visualization figures. NS and CTF performed the formal analysis of the structure and docking predictions of the antibodies. All authors wrote the original draft of the manuscript. All authors read and approved the final version of the manuscript.

## Acknowledgements

We gratefully acknowledge all GISAID data contributors (i.e., the authors and their originating laboratories) responsible for obtaining the specimens, and their submitting laboratories for generating the genetic sequence and metadata and sharing via the GISAID Initiative, on which this research is based.

We acknowledge the following entities at the University of North Carolina at Charlotte: Academic Affairs, The Office of Research, The Center for Computational Intelligence to Predict Health and Environmental Risks (CIPHER), The Department of Bioinformatics and Genomics, The College of Computing and Informatics, and the University Research Computing group. We gratefully acknowledge the support of the Belk Family.

## Data Sharing Statement

All code, data, results, and additional analyses are openly available on GitHub at: https://github.com/Santollan/Frankies. This repository includes the open-source logic for running the *Frankies* pipeline. Also, this includes all sequences and folded structures for the reference antibodies and H5N1 isolates used in this H5N1 study, analysis scripts, and docking metrics. This also includes the experimental outputs for the 30 diffused Fv candidates.

## Supplementary Materials

**Fig. 1.**
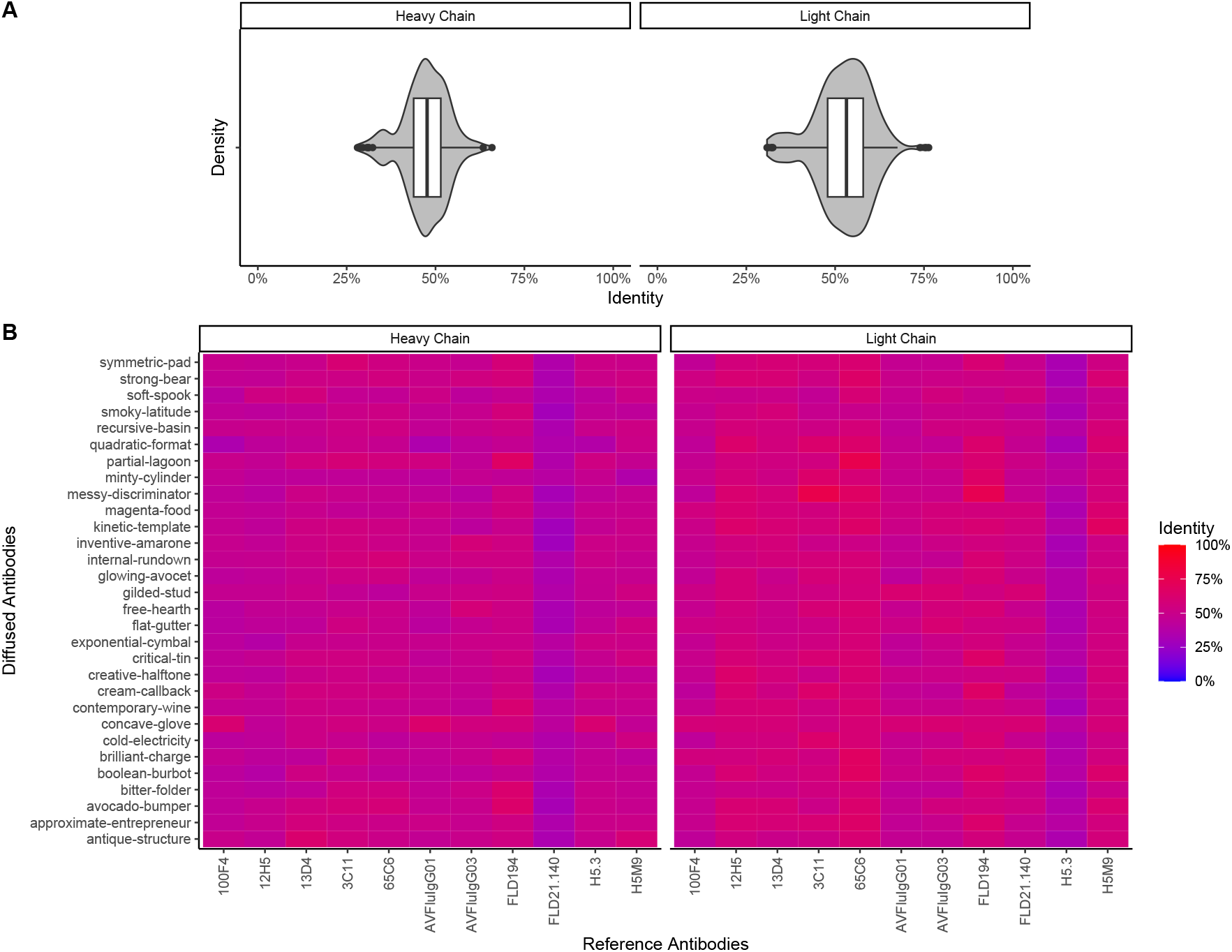
Sequence identity comparisons between the diffused and reference antibody sequences. A) Violin/box plots showing the distribution of the diffused sequences’ identities by chain. B) Heatmap showing the pairwise identity comparisons.

